# Weekly dance training over eight months reduces depression and correlates with fMRI brain signals in subcallosal cingulate gyrus (SCG) for people with Parkinson’s Disease: An observational study

**DOI:** 10.1101/2022.10.14.512180

**Authors:** Rebecca E Barnstaple, Karolina Bearss, Rachel J Bar, Joseph F DeSouza

## Abstract

Depression affects 280 million people globally and is considered a prodromal feature for increasingly prevalent neurodegenerative conditions including Parkinson’s disease (PD). With age-related neurodegeneration on the rise, it’s important to consider non-invasive, inexpensive interventions such as dance. Dance has emerged as a complementary treatment that may support adaptive neuroplasticity while diminishing motor and non-motor symptoms including depression. Although dance has been shown to impact brain structures and functions with improvements in motor and psychological symptoms, the neural mechanisms underlying depression/mood remain elusive. Our observational study tracks the relationship between depression scores and functional neuroimaging measures for subgenual cingulate gyrus (SCG). While learning choreography over an 8-month period, 34 dancers [23 people with PD] completed GDS questionnaires before and after their community dance classes. Seventeen of these dancers had BOLD fMRI scans conducted using learning-related protocols to examine underlying neural mechanisms for moving to music over 4 times points of learning. A significant decrease in depression scores correlated with a reduced BOLD signal from SCG, a putative node in the neural network of depression. Conclusions: This is the first study to clearly find a correlation with a neural substrate involved in mood changes as a function of dance for PD. Not only do the results contribute to understanding neural mechanisms involved in adaptive plasticity with a learning task, but they also uncover reduced activity within SCG during longitudinal therapeutic dance interventions. These results are especially illuminating since SCG is a controversial target in deep brain stimulation (DBS) used in the treatment of major depressive disorder (MDD).

## Introduction

Neurodegenerative conditions affect an increasingly larger proportion of the population with more than 10 million people worldwide living with Parkinson’s Disease (PD). It is the second most common neurodegenerative condition following Alzheimer’s and related dementias, and the fastest growing neurological condition (GBD 2013 Risk Factors Collaborators, 2015). Most commonly reported PD symptoms pertain to movement and nonmotor symptoms (NMS) such as anxiety, depression, fatigue, apathy, cognitive disturbances and dementia (Weintraub et al 2011; Bastide et al., 2015; Parkinson, 2002). It significantly impacts health and quality of life (QoL), with recent evidence suggesting that depression may be premorbid to the development of PD. Anxiety disorders, which frequently present prior to or in conjunction with major depression (Fava et al., 2000), have been shown to be indicative of prodromal stages of PD (Savica et al., 2010) and are also associated with disease progression (Dallé & Mabandla, 2018).

Despite the high incidence and impact of depression on QoL, the most prevalent treatments for PD - dopamine replacement therapy and deep brain stimulation (DBS) - target motor symptoms. Traditional pharmacological interventions for PD such as Levodopa and related medications can also be accompanied by unpleasant side effects, with efficacy subject to decay over time. Furthermore, surgical intervention with DBS is only available to suitable candidates, is not always successful, and may exacerbate some symptoms (Abbes et al., 2018).

Over the past two decades, a growing body of research has indicated that there are extensive benefits for people with PD (PwPD) associated with engaging in dance, a low-cost and readily available activity with historic and cultural precedents in many cultures and communities. Dance-based programs specifically designed for this population have shown to reduce motor impairments through improving balance and decreasing motor symptom severity, while also being linked with positive changes in nonmotor symptoms (NMS) such as mood (Hackney & Earhart, 2010; Hashimoto et al., 2015; Koch et al., 2016; Fontanesi & DeSouza, 2020; Bearss & DeSouza, 2021) and QoL (Westheimer, 2008; Bearss et al., 2017; Kalyani et al., 2019).

Dance for PD^®^(DfPD^®^), a dance program designed for PwPD and used internationally, is open to participants of all ages and stages of disease. In the DfPD^®^model, dance instructors are trained to ensure classes are appropriate and beneficial for PwPD. PwPD are invited to explore movements based in ballet and contemporary dance which are adapted to a range of motor abilities. While emphasising artistry, creativity, and beauty, DfPD maintains the central importance of dance by treating participants not as patients, but as dancers; DfPD also provides a respite from PD diagnosis and an alternate means of experiencing the body (Fontanesi & DeSouza, 2020; Houston & McGill, 2013; Houston, 2015; Westheimer, 2008; Westheimer et al., 2015).

To date, there has been only one fMRI case study with a single participant in which correlations between motor improvements and neural changes were explored. The authors found significant improvements on the Fullerton Advanced Balance scale and putative brain network connectivity between the basal ganglia and premotor cortices (Batson et al., 2014). In our recent study, a significant correlation between the levels of General Positive Affect (GPA) after DfPD and average Skin Conductance Levels (SCL) during a dance session was found, with significantly higher SCL rates during DfPD compared to a matched-intensity exercise; this suggests specific and additional benefits for DfPD (Fontanesi & DeSouza, 2021). While many studies of dance for PwPD have shown promising behavioral results, they are often limited by small sample sizes and a brief intervention period (weeks to a few months). Additionally, the time course and duration of these effects are unknown, particularly for NMS such as depression.

The present observational study examined the effects of attending weekly dance for PD classes on NMS associated with depression scores over 8 months at 4 sampling times. We hypothesized that multisensory learning associated with engaging in dance may act as a buffer against neurodegenerative disease (Bearss & DeSouza, 2021), and that this would be visible through modulations in brain signals over time. Specifically, we looked at changes in brain signals from an important, and still controversial node within the depression network (SCG), which showed correlations to reduced depression scores pre and post dance class.

## Methods

### Behavioural dance training and performance schedule over 8-months

All participants engaged in weekly 75-minute community-based Dance for PD^®^(DfPD^®^) structured dance classes which begins with a seated component, followed by mirroring and paired exercises, and ends with standing, while executing choreographed sequences and locomoting through space (see Table 1 from Bearss et al., 2017). Participants learned-novel choreography and practiced it at the end of each weekly class culminating in a public presentation on three separate occasions within the year of MRI scanning: Sharing Dance Day, Toronto City Hall, and at a Parkinson’s Central meeting.

### Study population – Behavioural testing

A total of 34 participants volunteered from the DfPD at Canada’s National Ballet School: 23 PwPD and 11 healthy controls (HC). PwPD were between the ages of 52-76 (*M*_age_ = 67.78, *SD* = 6.14; *n_Males_* = 9; *M_DxDuration_*= 5.56 years, range = 0–17 years). HC were between the ages of 61-83 (*M*_age_= 70.11; *SD* = 7.4; *n_Males_*= 6). Participants completed the Geriatric Depression Scale (GDS) before and after a single dance class (1.25 hours) at three separate time points: March, April and June. Motor symptoms were also examined at three time points using the Berg Balance Scale (BBS) and the Timed Up and Go (TUG) measures, as reported in Bearss et al. (2017).

### Study population – Neuroimaging sessions over 8-months

10 PwPD (ages 52-76 [*M*_age_ = 67.78, *SD* = 6.14]; *n_Males_* = 9; *M_DxDuration_*= 5.56 years, range = 0–17 years) completed a minimum of 2 of 4 functional imaging sessions in September, December, January and April at York University’s Sherman Health Sciences Centre to be included in further analyses (4 PwPD and 3 controls completed only one MRI scan). Disease progression ranged from asymptomatic to severe, as measured by the Hoehn and Yahr (H&Y) (0–4; *M_H&Y_* = 0.8). Written informed consent was obtained using an approved protocol from York University’s Ethics Board (2013-211 & 2017-296).

### Procedures - Imaging

A 3T Siemens Tim Trio MRI scanner was used to acquire functional and anatomical images using a 32-channel head coil. T2*-weighted echo planar imaging using parallel imaging

(GRAPPA) with an acceleration factor of 2X with the following parameters: 32 slices, 56 × 70 matrix, 210 mm × 168 mm FOV, 3 × 3 x 4 mm slice thick, TE = 30ms, flip angle of 90°, volume acquisition time of 2.0 s, for a total of 240 volumes per scan. Echo-planar images were coregistered with the high-resolution (1 mm^3^) anatomical scan of the subject’s brain taken at the end of each session (spin echo, TR = 1900ms, TE = 2.52 ms, flip angle = 9°, 256 x 256 matrix). Each subject’s head was restrained with padded cushions to reduce head movement artefacts.

While in the MRI, subjects were instructed to visualize themselves dancing from an internal first-person perspective while listening to music associated with the choreography learned in class (for more detail on this protocol see Bar & DeSouza, 2016). The choreography was a total of 3-minutes long, however, only the first minute of the music (A. Copeland’s - Hoe-down) was used during the fMRI scanning since they all had the most experience with this section. The dance visualization task employed a blocked design 30 s *OFF* and 60 s *ON; ON* states were alternated five times with 30-s periods of rest for a total scan time of 8-min. The anatomical seed region was set using landmarks from Mayberg et al. (2005) and Hamani et al. (2011) with a surrounding 5-cm radius in BrainVoyager 22.0 (v22.0.3.4578, Maastricht, The Netherlands). Following statistical analysis of the BOLD signal, data was conducted in MATLAB (The MathWorks Inc., version 9.8.0.1417392, R2020a) and RStudio (version 1.2.5033).

### Preprocessing

Functional scans were superimposed on anatomical brain images, aligned on the anterior commissure-posterior commissure line, and transformed into Talairach space in BrainVoyager 22.0. Functional data from each scan were screened for motion and/or magnet artifacts to detect abrupt movements of the head. In addition, we ensured that no obvious motion artifacts were present in the activation maps from individual PwPD. Participants were video recorded while in the fMRI scanner, providing a record of whether any noticeable physical movements occurred during scanning. None of the 28-functional scans from 10 PwPD were removed due to motion artifacts.

### Statistical Analysis

We used an anatomically defined seed region (radius 5mm) corresponding with the coordinates of SCG as modulation of activity in this area has been associated with changes in depression and mood (Mayberg, 2005; Hamani et al., 2011). Functional data were analyzed for changes during music-visualisation task periods over the four time points collected.

## Results

### Behavioural - Geriatric Depression Scale - GDS

Data from the GDS were analyzed in SPSS statistical software (version 24.0; IBM Corp 2016) employing a linear mixed effects model analysis to account for individual subject data variability and allow for inclusion of data from participants missing one or more data collection points across time (March, April, June).

Statistical analyses showed improvements in depression/mood scores when examining for the effects of the dance class over time. There was a significant main effect for time [*F* (4, 135.31) = 3.677, *p*< 0.01] and the measured effect of the dance class on GDS (pre/post) [*F* (1, 125.97) = 5.266,*p* < 0.025]; no significant interaction was found between experience (time in months dancing) and the GDS impact following a 75-minute dance class [*F* (2, 125.85) = 0.166, *p* = 0.848].

Figure 1 shows a significant reduction of GDS scores across time pre/post dance class for March and April, plus pre/post the dance class for the months of data collection. Further examination of the pre/post class comparisons used paired *t*-tests to examine the significance of the conditions (pre/post) across the differing time points and found a significant reduction in reported GDS symptoms. Significant differences in GDS scores were observed within the PwPD group when examining GDS scores for the pre/post class at each of the time points as well as looking at the condition between specific time points. A previous publication reports on motor improvements (Bearss et al., 2017); BBS and TUG analysis showed significant improvements for PwPD.

**Figure 1:**
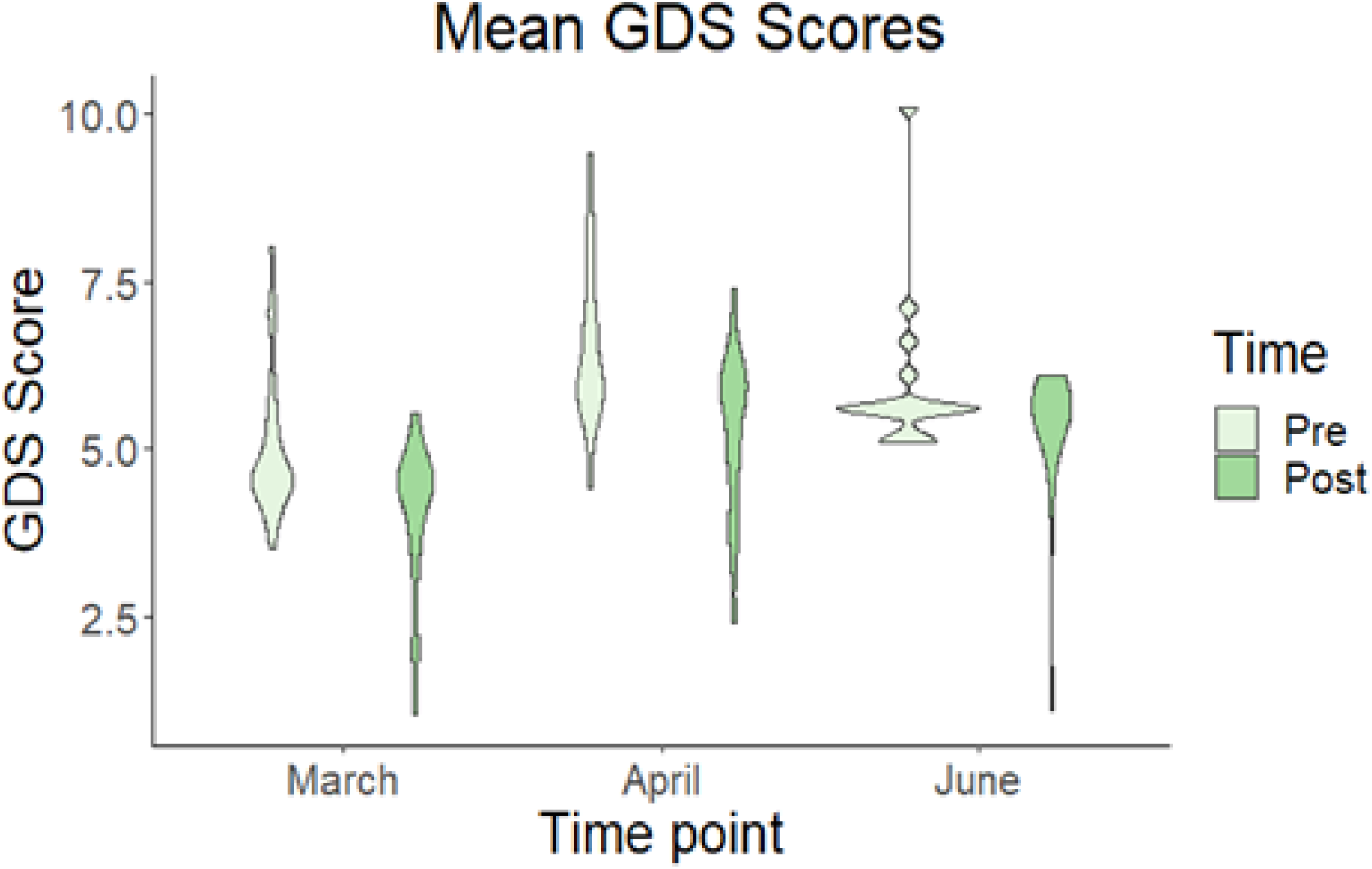
Mean GDS scores across all dance participants before (PRE) and after (POST) a dance class conducted in the dance studio in March, April and June.

### Functional Imaging anatomical seed location

Based on literature showing this to be a crucial node in affective networks associated with Major Depression Disorder (MDD), an anatomical seed localizer for SCG (Figure 2A) was used in extracting BOLD signals for this analysis. Figure 2B shows GDS scores for participants who were scanned in the MRI at the appropriate times in relation to the dance class. Figure 2C shows the averaged signal across subjects for the one minute of music where the PwPD visualized their learned dance at four different times in learning (September, December, January and April).

**Figure 2:**
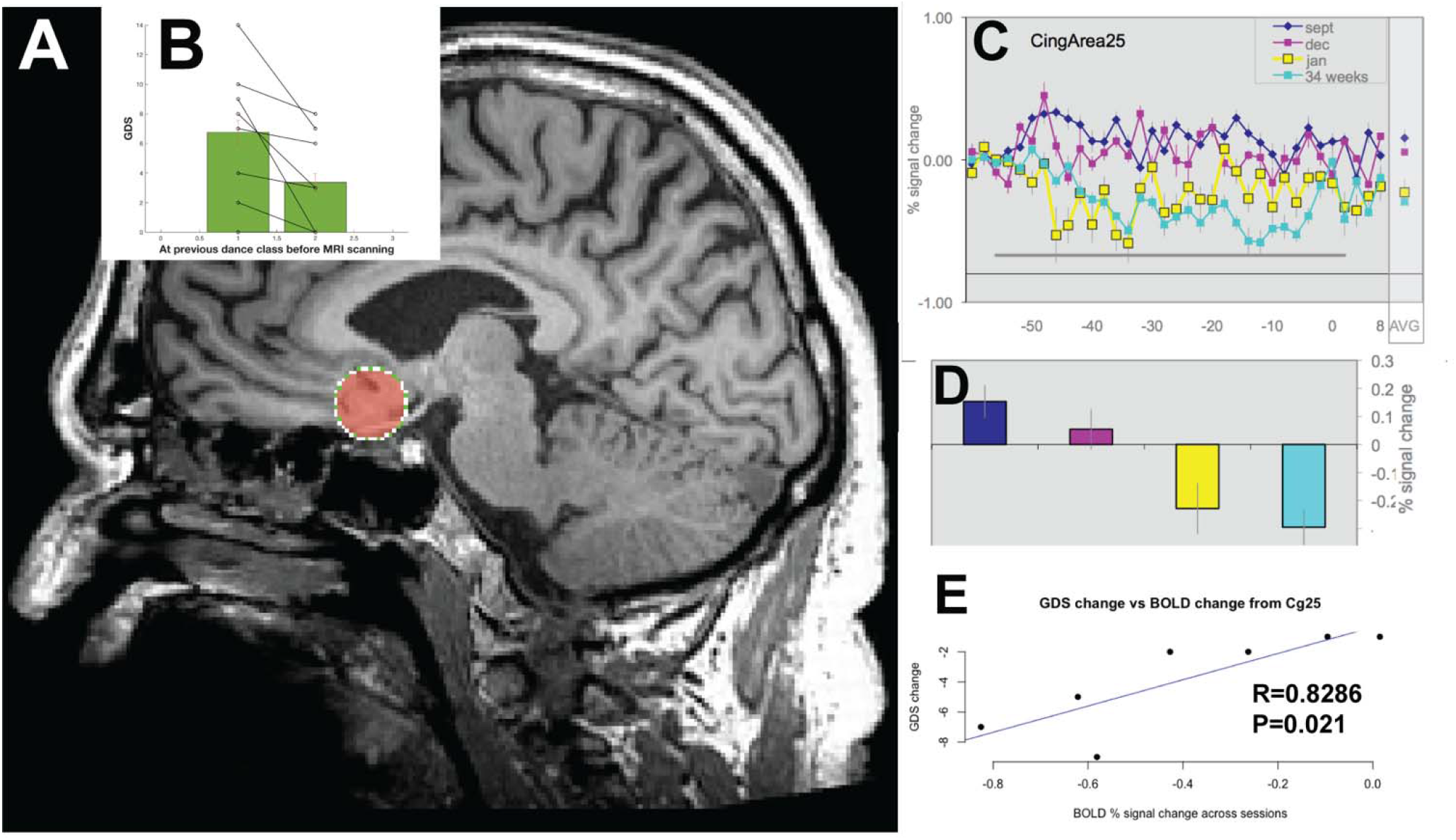
**A.** Anatomical location of seed in Subcallosal Cingulate Gyrus (SCG) to probe BOLD signals while people with Parkinson’s disease visualized learned choreography elicited by music trained in the DfPD class. **B**. Geriatric Depression Scores (GDS) for this subset of PwPD <u>(p<0.05, _paired t-test,</u> n=7 PwPD). **C**. BOLD signals averaged across PwPD for the music visualization training task during September, December, January and April for all 10 PwPD (28-scans). Data aligned to the end of the one minute of music. Last point is the average plot across the grey bar for the one minute of music visualization. **D**. Bar graphs showing averaged BOLD signals from SCG with standard error of the mean across PwPD. **E**. A significant correlation for the change in GDS scores (from Fig B) and BOLD signals reductions (from Fig D) for the seven PwPD who both were scanned in the MRI and completed the GDS questionnaires at the dance studio.

There was a decrease in BOLD signal from area SCG as the months progressed at all time points (Figure 2D). A Pearson correlation analysis of the change in GDS data from pre to post (Figure 2B) and the decrease in BOLD signal data showed a strong significant positive correlation (*r*(5)= 0.83, *t*=-3.31,*p* <0.025) (Figure 2E).

## Discussion

Depression, in its many potential manifestations, can be a serious and debilitating disorder impacting every aspect of life. It’s also incredibly common; in 2020, an estimated 6% of all U.S adults suffered at least one major depressive episode (NSDUH, 2021). Despite the prevalence of depression, the neural mechanisms underlying its development, expression, and treatment remain poorly understood. As an example of the complexity of this subject, the commonly held hypothesis that low serotonin levels are linked with depression is now under debate, emphasizing the need to develop and further investigate novel treatments (Moncrieff et al., 2022). Our research contributes initial evidence that participating in a social activity such as dance can alter activity in a brain region associated with depression/mood, and thus may further this project.

Longitudinal reductions in BOLD brain signal from SCG, an area believed to be a critical node in a corticolimbic network associated with depression (Hamani et al., 2011), strongly correlated with decreases in depression scores (GDS) during our dance program in PwPD. This observational study combines local data collected at the intervention site (dance studio) with functional neuroimaging. In the MRI scanner, participants mentally rehearsed learned movement patterns associated with their in-studio choreographed dance sequences, while listening to the music that accompanies the dance. A reduction in BOLD activation in a region involved in depression was found. This provides preliminary evidence that participation in dance classes involving learned choreography can modulate neural activity in a way that is adaptive.

Many imaging studies suggest increased activity in the SCG region for patients with depression (Mayberg et al., 2005; Hamani et al., 2011), and reductions in this activity following treatment with antidepressants, placebos, repetitive TMS (transcranial magnetic stimulation), electroconvulsive therapy, deep brain stimulation and cingulotomy (Hamani et al., 2009; 2011). However, none of these studies were conducted with people with PD or used dance as the intervention. Our investigation provides initial evidence that brain signals from SCG decrease in PwPD who participate in noninvasive weekly community dance classes over eight months.

SCG plays an important role in regulating emotion. Located in the subgenual anterior cingulate cortex (subgenual ACC), degeneration in this area correlates with depressed mood and anhedonia (Rudebeck et al., 2014). Specifically, SCG is thought to be a critical node in regulating a corticolimbic network with increased activity in this area associated with depression (Mayberg et al., 2005). Also, SCG is a primary target in DBS for treatment-resistant depression (TRD) (Roet et al., 2020). It is hypothesized that DBS can modulate the depression network through a reduction of SCG activity (Mayberg et al., 2005; Hamani et al., 2011), leading to antidepressant effects.

Depression is one of the most prevalent and debilitating nonmotor symptoms faced by people living with PD and may even precede motor disturbances. It is also one of the most challenging to identify and treat because psychomotor slowing and blunted affect (masking) can contribute to missed diagnosis of depression. Importantly, interactions between some antidepressants (SSRIs) and PD medications, in some cases, may potentially exacerbate motor symptoms. With increased prevalence of depression across a wide variety of populations, including PwPD, new treatments and models are sorely needed.

In the wake of COVID-19, there has been a global increase in mental health concerns closely associated with reduced access to social activities and emotional support. Group movement experiences associated with health, art, and ritual have played a role in human culture for millennia. Beyond their aesthetic value, it is entirely possible that they may play an important role in regulating mood and adaptive behaviours. Our participants showed a consistent decrease in BOLD signal activity in SCG over an eighth month period, while participating in weekly group dance sessions that involved learning a new motor sequence set to music, an experience that was re-elicited in the MRI scanner.

Our match between biological (fMRI) and behavioural measures (GDS) provides early evidence that participation in a program such as Dance for PD, a non-invasive, widely available intervention, can facilitate adaptive plastic changes in neural networks associated with depression. This is an important finding in that it demonstrates how socio-cultural experiences may be integrated into or may affect biological circuits. These initial preliminary observational results require further investigation, ideally through a randomized control trial (RCT) to better elucidate the contribution of complex neural mechanisms and reduce the potential confounds that accompany any observational study.

## Acknowledgements

Thanks to all the people from our lab over the years for the project and all the people at our testing locations. We are indebted to S. Robichaud, D. Rabinovich, R. Cohan, P. Dhami, S. Maguire, H. Tehrani, K. McDonald, and R. Andrew for invaluable assistance during all phases of the research program. A special thanks to Dr. J. Bek and P. Snider for comments on this manuscript, S. Houshangi-Tabrizi for figure composition and all the lab members (**http://www.joeLAB.com**) for their dedication since study inception. We thank S. Ciantar for compiling, writing and running the statistics in SPSS for the GDS. We are also thankful for the collaborations with Canada’s National Ballet School (A. Seto, A. Powell) and Dance with Parkinson’s Canada (S. Robichaud) and Trinity St. Paul’s Church in Toronto, ON, Canada and D. Leventhal from Dance for Parkinson’s at Mark Morris Dance Group for continued training and support. Author contributions: JFXD conceived, executed, collected, analyzed, contributed resources, project administration, funding acquisition and edited the writing. RJB conceived, executed, funding acquisition and contributed to writing. KB collected, analyzed, and contributed to writing. REB conceived, analyzed, and contributed to writing. All authors have read and agreed to the published version of the manuscript. Funding: Parkinson Canada Pilot grant to JFXD; JFXD funded by National Science and Engineering Research Council (NSERC) Discovery Grant (2017-05647) and donations from the Irpinia Club of Toronto and others to JFXD.

